# ADHESION, BIOFILM, AND INVASION: INVESTIGATING THE VIRULENCE MECHANISMS OF *Prevotella* spp.

**DOI:** 10.64898/2026.01.15.699568

**Authors:** A.T.O Marre, J.C.C. Morgado, S.B Costa, I.S Barcellos, V.C.A.C Chuva, A.C.S.C Oliveira, L.A. Lobo

**Affiliations:** Federal University of Rio de Janeiro; Rio de Janeiro State University; Universidade Federal do Rio de Janeiro (UFRJ)

## Abstract

The *Prevotella* genus are strict anaerobic organisms associated with opportunistic infections in the vaginal, oral, and gastrointestinal cavities. During infection, virulence mechanisms such as adhesion to host tissues, invasion of cells and connective tissue, and evasion of the immune system are essential for bacterial establishment and host persistence. In the present study, we investigated the adhesion to human extracellular matrix proteins, biofilm formation, Matrigel invasion, and plasminogen activation of strains from the Prevotella species, including *P. intermedia*, *P. melaninogenica*, and *P. nigrescens*. The bacterial adhesion capacity was quantified by the interaction of these bacteria with extracellular matrix proteins, including fibronectin, collagen type IV, collagen type I, laminin type 1, and Matrigel, previously immobilized on glass slides. *P. intermedia* and *P. nigrescens* demonstrated adhesion to fibronectin, type IV collagen, and Matrigel. *P. melaninogenica* did not adhere to the substrates under the study conditions. To identify ligands in *Prevotella* and *Fusobacterium*, outer membrane protein extracts were purified from *P. intermedia*, *P. nigrescens*, and *F. nucleatum* and subjected to affinity chromatography using NHS-activated Sepharose columns containing immobilized laminin, fibronectin, and type IV and type I collagen. Eluted fractions containing potential ligands were sent for mass spectrometry analysis. In *P. intermedia*, six proteins were identified as potential laminin adhesins and 15 as potential type IV collagen adhesins. In *P. nigrescens*, five proteins were identified as potential laminin adhesins and three as potential type IV collagen adhesins. Biofilm experiments were also conducted in the presence and absence of Matrigel. Biofilm formation was reduced in the presence of this substrate in *P. intermedia* and *P. melaninogenica*, while no significant difference was observed in the other species tested. To analyze the biofilm architecture, scanning electron microscopy was performed. It was observed that, in the presence of Matrigel, the biofilm surface of the analyzed species was altered. *P. melaninogenica* did not form biofilm on the glass surface used for SEM. A transwell invasion assay was performed for all *Prevotella* species. It was observed that only *P. melaninogenica* was capable of crossing the Matrigel layer. Matrigel degradation assays using SDS-PAGE showed that *P. melaninogenica* degrade matrix proteins and plasminogen. To understand the interaction between species and plasminogen, a plasminogen activation kinetic assay was conducted, in which only *P. melaninogenica* activated this molecule, likely utilizing this strategy to destroy tissues. Understanding the mechanisms involved in virulence may help develop new strategies to prevent periodontitis and biofilm formation in the gingival sulcus.

## Introduction

Adhesion is considered a crucial step in the establishment of infection, as adherent bacteria are not removed by cleansing mechanisms such as peristalsis and ciliary movement, nor by immune system responses. Bacterial adhesion to host surfaces confers several advantages, including strong attachment and resistance to removal by hydrodynamic shear forces [1;2] access to nutrients released by host tissues [3], and protection against the deleterious effects of antimicrobial agents [4;5;6]. Components of the human extracellular matrix (ECM) are potential targets for the adhesion of pathogenic bacteria [7;8;9]. These ECM molecules can be exposed through mucosal injuries acquired in various ways, such as brushing, cuts, or epithelial disruption. Numerous bacterial proteins can recognize and bind to one or more ECM components [10; 5]. Owing to their diversity, these microbial structures are collectively known as **Microbial Surface Components Recognizing Adhesive Matrix Molecules (MSCRAMMs)** [10; 11; 5].

Periodontal diseases are characterized as a group of inflammatory conditions affecting the gingiva, bone, and connective tissues that support the teeth [12]. Gingivitis and chronic periodontitis are initiated and sustained by the activity of microorganisms present in the dental biofilm [13]. During the formation of this biofilm, microorganisms must firmly adhere to surfaces, since they can be removed by saliva and food. Genera such as *Prevotella*, *Porphyromonas*, *Treponema*, *Aggregatibacter*, and *Eubacterium* spp. are commonly found in the biofilm, being the bacteria most closely associated with oral infections [14].

Species of the genus Prevotella contribute to ecological shifts that promote biofilm maturation and progression toward periodontal pathology. Although they are associated with disease, under conditions of immune homeostasis they act primarily as commensal members of the oral microbiota. Several clinically important species—*P. intermedia*, *P. nigrescens*, *P. denticola*, *P. disiens* and *P. melaninogenica*—have been isolated from the subgingival plaque of patients with periodontitis [15; 16]. Within the ecological model of periodontal disease, these species are considered common periodontal pathogens and are categorized within the orange microbial complex [17]. Evidence further suggests that Prevotella species contribute to disease development through interactions with members of other pathogenic consortia. For example, Kamaguch and colleagues proposed that *P. intermedia* facilitates the establishment of *Porphyromonas gingivalis*, a key member of the red complex, within dental plaque [18]. This cooperative relationship highlights the ecological and functional significance of *P. intermedia* in oral biofilm maturation and periodontal pathogenesis. Similarly, *P. nigrescens*, also assigned to the orange complex, is regarded as a clinically relevant species strongly associated with periodontal disease [19].

Given the prevalence of *Prevotella* across multiple infectious scenarios and its broad repertoire of pathogenic mechanisms, deeper investigation into the virulence traits of this genus is warranted. Particular attention should be directed toward species most frequently isolated from disease-associated sites—including *P. intermedia*, *P. melaninogenica*, and *P. nigrescens*—as these taxa likely represent central contributors to the dysbiotic transitions underlying periodontal pathology.

## Materials and methods

### Bacterial culture conditions

*Prevotella intermedia* ATCC 49046, *Prevotella melaninogenica* ATCC 25845, *Prevotella nigrescens* ATCC 33563 (American Type Culture Collection) and *Staphylococcus epidermidis* ATCC 35984 stocks were maintained frozen in -80°C in a 20% glycerol solution and were routinely cultivated on 5% blood agar supplemented with hemin (0.5 mg/mL) and menadione (1 mg/mL) under anaerobic conditions with 80% N2, 10% H2, and 10% CO2 at 37°C.

### Adhesion assay

ECM proteins, type IV collagen, fibronectin and laminin, at a concentration of 0.02 mg/mL [20], were immobilized on glass slides for 24 hours at 25°C. After protein adsorption, the plate is blocked for 1 hour with the blocking solution (1% BSA, 1% gelatin, and 0.1% Tween 20) to prevent nonspecific interactions. For the interaction, colonies of *Prevotella* species grown on blood agar were suspended in phosphate-buffered saline (PBS) and adjusted to a concentration of 5 x 10^7 CFU/mL using a spectrophotometer (OD 550 nm). The bacterial suspension was added to the slides and incubated for 1 hour at 37°C in anaerobic conditions. After the interaction, the plate is washed three times with cold PBS and fixed with 3.7% paraformaldehyde for 15 minutes. The slides are removed from the plate and placed on a microscope slide for analysis.

### Statistics analysis

All data obtained from the adhesion quantification experiments were subjected to statistical analyses using Graphpad Prism 8 software. A test was conducted to confirm whether the samples were normally distributed, utilizing the Normality and Lognormality tool. After confirmation, analysis of variance (ANOVA) was performed. The ANOVA analyses were unpaired (independent samples) and non-parametric (not normally distributed). A p-value of <0.05 was considered significant for results with a meaningful difference. Following the initial ANOVA analysis, a Tunn’s test was conducted to compare two groups of data with the initial ANOVA analysis, considering multiple comparisons among the data in relation to the main analysis.

### OMP extraction

The strains of *P. intermedia*, *P. melaninogenica*, and *P. nigrescens* were grown in BHI broth for 16 hours at 37°C under anaerobic conditions. After growth, the broth was subjected to centrifugation at 8000 xg for 20 minutes. The pellets were resuspended in Hepes buffer (10 mM, pH 7.4 - Sigma®). The cells in the pellet were disrupted using a Microson™ Ultrasonic Cell Disruptor sonicator (NY, USA) at 40 Watts for 10 cycles of 90 seconds each. Separation of intact and disrupted cells was performed by centrifugation at 4000 xg for 10 minutes. The supernatant containing the envelope fraction was subjected to ultracentrifugation (Beckman Coulter Inc., California, USA) at 100,000 × g for 1 hour using an SW40Ti rotor. The resulting pellet was resuspended in HEPES buffer (10 mM, pH 7.4) and ultracentrifuged again under the same conditions. After 1 hour, the pellet was resuspended in 1% N-lauroylsarcosine (Sarkosyl, Sigma®) prepared in HEPES buffer (10 mM, pH 7.4) and incubated at 37°C for 60 minutes with agitation. The N-lauroylsarcosine-treated membrane proteins were subjected to a third ultracentrifugation under the same conditions described above. This step yielded an extract enriched in PMEs. The pellet was washed with HEPES buffer and ultracentrifuged once more under the same conditions. Following the final centrifugation, the pellet was resuspended in HEPES buffer, supplemented with 10 µL of RapiGest (0.2%), and stored at −20 °C.

### Affinity Chromatography for ligand isolation

The enriched extract of outer membrane proteins was subjected to affinity chromatography using immobilized laminin and collagen type IV for the purification of the specific ligand. In this experiment, the HiTrap™ NHS-activated HP column (GE Healthcare Life Sciences) was used, which is activated sepharose for coupling ligands containing primary amino groups. The experiment was conducted following the protocol provided by the manufacturer. All extracts that were eluted from the column were stored at -20°C.

### *In solution* protein digestion

A concentration of 0.01 µg/µL of the purified protein was used for the experiment. Ammonium bicarbonate (100 mM, pH 8.5) and calcium chloride (10 mM) were added to the protein sample, followed by the addition of 0.2% RapiGest. The mixture was incubated at 80 °C for 15 minutes in a thermocycler, then centrifuged at 6,000 × g for 5 minutes. Subsequently, dithiothreitol (100 mM) was added and the sample was incubated at 60 °C for 30 minutes. Iodoacetamide (300 mM) was then added, and the solution was kept in the dark for 30 minutes. A trypsin solution (Promega), prepared in 10 mM ammonium bicarbonate with 1 mM CaCl₂ (pH 10), was added to the sample, which was then incubated overnight at 37 °C in a thermocycler. After digestion, trifluoroacetic acid was added and the sample was incubated at 37 °C for 90 minutes. The reaction mixture was subsequently centrifuged at 16,000 × g for 30 minutes at 6 °C. The supernatant was transferred to a new tube and dried in a SpeedVac. Finally, the samples were stored at −20 °C for mass spectrometry analysis.

### Peptide mass fingerprinting

The protein samples were resuspended in a solution of 0.1% formic acid for analysis by liquid chromatography coupled to mass spectrometry (LC-MS) in a NanoElute 2 system (Bruker) coupled to a Maxis Impact. The column model employed was the PepSep C18, 15 cm x 75 µM, with 1.9 µM particles, along with a Thermo Fisher C18 Trap column, 5 cm x 3 mm. The run was performed using a gradient of mobile phase A (water with 0.1% formic acid, v/v) and mobile phase B (acetonitrile with 0.1% formic acid, v/v). The analysis and identification of the peptides were performed using an ESI-Q-TOF (Bruker), with a nanoelectrospray ionization source and positive polarity. The method for acquiring precursor and peptide fragment spectra was data-dependent acquisition (DDA/AutoMs), with fragmentation of 20 precursors per cycle.

The analysis of the generated mass spectra was performed using MaxQuant software (v2.4.9.0), utilizing the National Center for Biotechnology Information (NCBI) database, considering Oxidation (M) and Acetyl (N-terminal protein) as variable modifications and Carbamidomethyl (C) as a fixed modification. The data generated by the program were cleaned by removing contaminants, decoy hits, and proteins with an MS/MS count of zero. Proteins with null label-free quantification (LFQ) intensity were also removed. The resulting proteins were submitted to the Cello program (v2.5) and the platforms AlphaFold, InterPro, NCBI, and UniProt.

The LFQ intensity values were exported and processed in Python 3.12 using the pandas (v2.2.0), numpy (v2.3.3), matplotlib (v3.10.7), and seaborn (v0.13.2) libraries. For each protein group identified by MaxQuant, the first Protein ID was used as the representative identifier for the group. LFQ intensity values equal to zero were treated as missing data and replaced with NaN (not a number). Data normalization was performed in two steps: first, LFQ intensity values were log₂-transformed to stabilize variance. Next, z-score normalization was applied independently for each protein (row-wise normalization), calculated as z = (x – μ) / σ, where x represents the log₂ intensity value, μ is the mean log₂ value for the protein across all samples, and σ is the corresponding standard deviation. This approach allows comparison of relative expression patterns across samples, independent of absolute differences in protein abundance. For co-occurrence analysis, a protein was considered present in an experimental group (collagen or laminin) if it was detected in at least one biological replicate (LFQ value > 0). Proteins were classified into three categories: collagen-exclusive, laminin-exclusive, or shared (present in both groups). Data visualization was performed using a hierarchical heatmap with proteins ordered alphabetically and a Venn diagram to represent co-occurrence relationships. The heatmap used a diverging colormap (coolwarm) centered at zero, where positive z-score values indicate intensities above the protein’s mean and negative values indicate intensities below the mean.

### Biofilm assay

To understand biofilm formation in the presence of extracellular matrix (ECM) components, biofilm assays were conducted with and without Matrigel (Geltrex, Corning). A 24-well plate was prepared with Matrigel at a concentration of 1:10 and then incubated at 37°C for 40 minutes to promote the polymerization of the ECM protein solution. After this period, inocula of *Prevotella* and *Staphylococcus epidermidis* were prepared in BHI medium with 0.5% glucose and adjusted to a concentration of 5 × 10^6 CFU/mL. The analysis of biofilm surface formation was performed using scanning electron microscopy (SEM). After incubation, the plate was prepared for SEM at the Multiuser Microscopy Unit Padrón-Lins (UNIMICRO). The plate was fixed with a solution of 25% glutaraldehyde, 0.2M sodium cacodylate, and Milli-Q water for 1 hour. Samples were washed with 0.1M sodium cacodylate buffer, followed by post-fixation using a solution of 1% osmium tetroxide and 0.1M sodium cacodylate. For the dehydration step, increasing concentrations of ethanol (30%, 50%, 70%, 90%, and 100%) were used. After this step, the samples were dried with a solution of hexamethyldisiloxane mixed with ethanol and pure hexamethyldisiloxane. Finally, the samples were sputter-coated in a Leica EM SCD 050 (Leica Microsystems, Germany) for 70 seconds at 40 mA. The samples will be analyzed at the Advanced Microscopy Unit of the National Center for Structure and Bioimaging (CENABIO) using a Zeiss EVO MA 10 microscope (Carl Zeiss, Göttingen, Germany).

### Transwell invasion assay

The experiment was conducted using Transwell inserts (Falcon®) with a 3.0 µM membrane. Matrigel (Geltrex™) was diluted in cold PBS at a ratio of 1:3 and added to each insert. The plate was incubated at 37°C for 4 hours to promote the polymerization of the Matrigel. The inoculum was prepared with *Prevotella* strains diluted in PBS to reach an optical density of 6 × 10^8 CFU/mL. The suspension was deposited on top of the polymerized Matrigel layer, and PBS was added to the well of the 24-well plate. The plate was then incubated at 37°C under anaerobic conditions. After 24, 48, and 72 hours, aliquots of the PBS in the well were removed and diluted to 1:100 in PBS. Subsequently, these were cultivated on blood agar at 37°C for 48 hours. All colonies that grew were counted and quantified.

## Results

### Adhesion test

To assess the adhesion capacity of oral *Prevotella* species, we first evaluated their interaction with extracellular matrix (ECM) proteins, including laminin, type IV collagen, and fibronectin. As shown in Fig. 1X, *P. intermedia* displayed robust adhesion to all ECM substrates tested. The mean adhesion values were 190.7 Bac/FC (bacteria per counted field) for laminin, 141.3 Bac/FC for fibronectin, 90.5 Bac/FC for type IV collagen, and 20.7 Bac/FC for the negative control (1% w/v BSA with 1% v/v Tween 20) (Fig. 1X). Adhesion of *P. nigrescens* to ECM molecules was also observed, as confirmed by microscopy (Fig. 1X). The mean adhesion to laminin was 166.1 Bac/FC, to fibronectin 132.8 Bac/FC, to type IV collagen 163.4 Bac/FC, and to the negative control 11.2 Bac/FC (Fig. 1X). In contrast, *P. melaninogenica* did not exhibit detectable adhesion to any of the ECM proteins under the experimental conditions

**Fig. 1.**
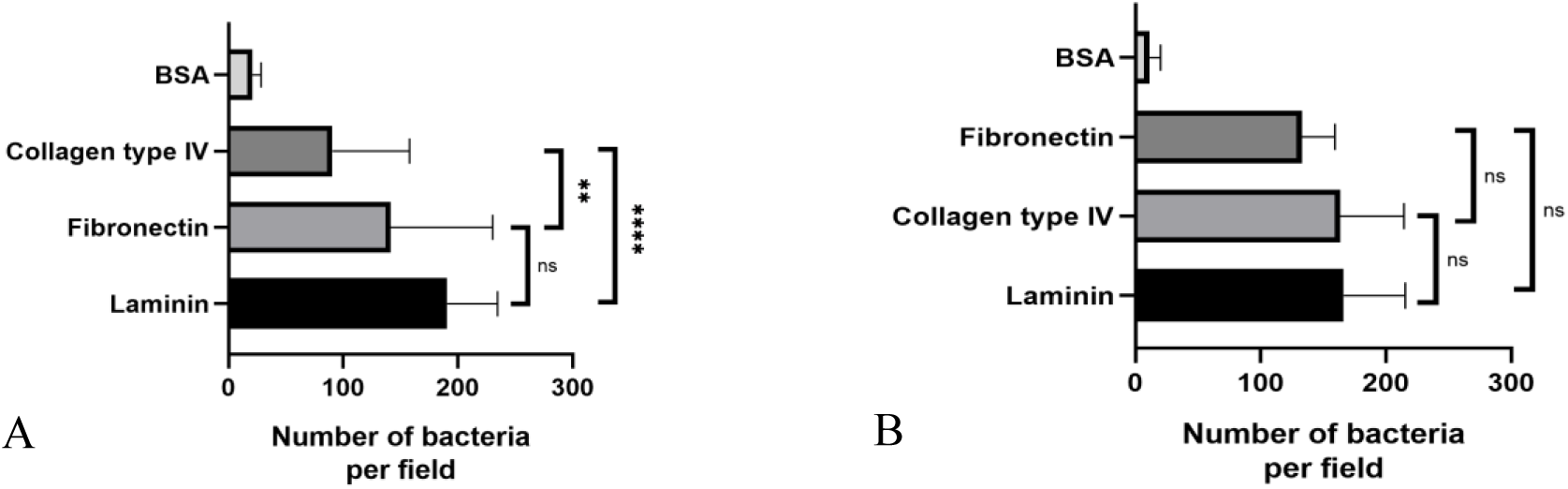
*Prevotella* species are capable of adhering to the MEC. (A) *Prevotella intermedia* quantification of adherence in glass coverslips. (B) *Prevotella* nigrescens quantification of adherence in glass coverslips. Bacterial counts were performed using ImageJ software, which quantifies particles in the image and provides a count of adhered bacteria per image. Each count was entered into GraphPad Prism 8, which calculates the mean adhesion and performs statistical analyses using one-way ANOVA followed by Tukey’s multiple comparisons test. The results presented represent the aggregation of two biological replicates and four technical replicates. Asterisks indicate statistical significance levels (****) – p < 0.0001, (**) - p = 0.0063, (ns) - p > 0.05. Error bars in each column represent the standard deviation.

### Analysis of the proteome of *P. intermedia* adhesion to collagen type IV and laminin

Following confirmation of the adhesion results, an affinity chromatography column using Sepharose beads was performed. The eluate containing the specifically bound fraction was subjected to analysis by nano-ESI-Q-TOF. Mass spectrometry analysis of the P. intermedia eluate revealed the presence of 15 proteins in the column enriched with type IV collagen (Table 1), whereas in the column enriched with type I laminin, the presence of 6 proteins was detected (Table 2). All identified proteins contained more than two unique peptides, which were exclusively associated with the identified protein group and were not found in other proteins. In addition, the protein groups did not present zero ion intensity, supporting that the identified peptides specifically belonged to the identified proteins. The proteins selected for bioinformatics analyses also presented LFQ values different from zero (Supplementary Table 1), and those with a Q-value ≤ 0.01 were considered reliable. Among the identified proteins, three were detected in both the laminin column analysis and the type IV collagen column analysis, namely WP_061869075.1, AFJ09446.1, and SUB95050.1. The presence of these proteins in both columns highlights their potential role as adhesins.

**Tabel 1:** *P. intermedia* collagen type IV.

| First_Protein_ID | Sequence coverage [%] | Score | Q-value | Unique peptides | Coverage_class | Confidence |
| --- | --- | --- | --- | --- | --- | --- |
| AFJ07558.1 | 181 | 67465 | 0 | 5 | High | High Fidelity |
| SUB95050.1 | 451 | 32331 | 0 | 6 | High | High Fidelity |
| AFJ09386.1 | 61 | 32331 | 0 | 3 | High | High Fidelity |
| AFJ09446.1 | 173 | 32331 | 0 | 12 | High | High Fidelity |
| APW31598.1 | 253 | 20423 | 0 | 6 | High | High Fidelity |
| APW31672.1 | 187 | 32331 | 0 | 5 | High | High Fidelity |
| WP_196755852.1 | 256 | 32331 | 0 | 9 | High | High Fidelity |
| APW31741.1 | 3 | 16106 | 0 | 2 | Low | Medium Fidelity |
| ATV29078.1 | 94 | 32331 | 0 | 3 | High | High Fidelity |
| PIK18545.1 | 197 | 32331 | 0 | 6 | High | High Fidelity |
| ATV54185.1 | 142 | 32331 | 0 | 12 | High | High Fidelity |
| WP_061868271.1 | 234 | 32331 | 0 | 3 | High | High Fidelity |
| WP_061868356.1 | 61 | 32331 | 0 | 2 | High | High Fidelity |
| WP_061869074.1 | 198 | 32331 | 0 | 10 | High | High Fidelity |
| WP_061869075.1 | 319 | 32331 | 0 | 32 | High | High Fidelity |

**Tabel 2:** *P. intermedia* laminin.

| First_Protein_ID | Sequence coverage [%] | Score | Q-value | Unique peptides | Coverage_class | Confidence |
| --- | --- | --- | --- | --- | --- | --- |
| AFJ07901.1 | 57 | 4453 | 0 | 3 | High | High Fidelity |
| SUB95050.1 | 463 | 32331 | 0 | 7 | High | High Fidelity |
| AFJ09446.1 | 205 | 14641 | 0 | 15 | High | High Fidelity |
| WP_198404996.1 | 194 | 32331 | 0 | 6 | High | High Fidelity |
| ATV54123.1 | 202 | 15532 | 0 | 7 | High | High Fidelity |
| WP_061869075.1 | 308 | 32331 | 0 | 31 | High | High Fidelity |

Heatmap analyses based on Z-score–normalized LFQ intensities revealed distinct patterns of protein abundance between the collagen and laminin conditions. The biological replicates showed high concordance, indicating good experimental reproducibility. Among the three recurring proteins, WP_061869075.1 exhibited above-average abundance when detected in the type IV collagen column and below-average abundance in the type I laminin column. The protein AFJ09446.1 displayed below-average abundance under both conditions, whereas SUB95050.1 showed above-average abundance in the type IV collagen condition (Figura 2).

**Figure 2:**
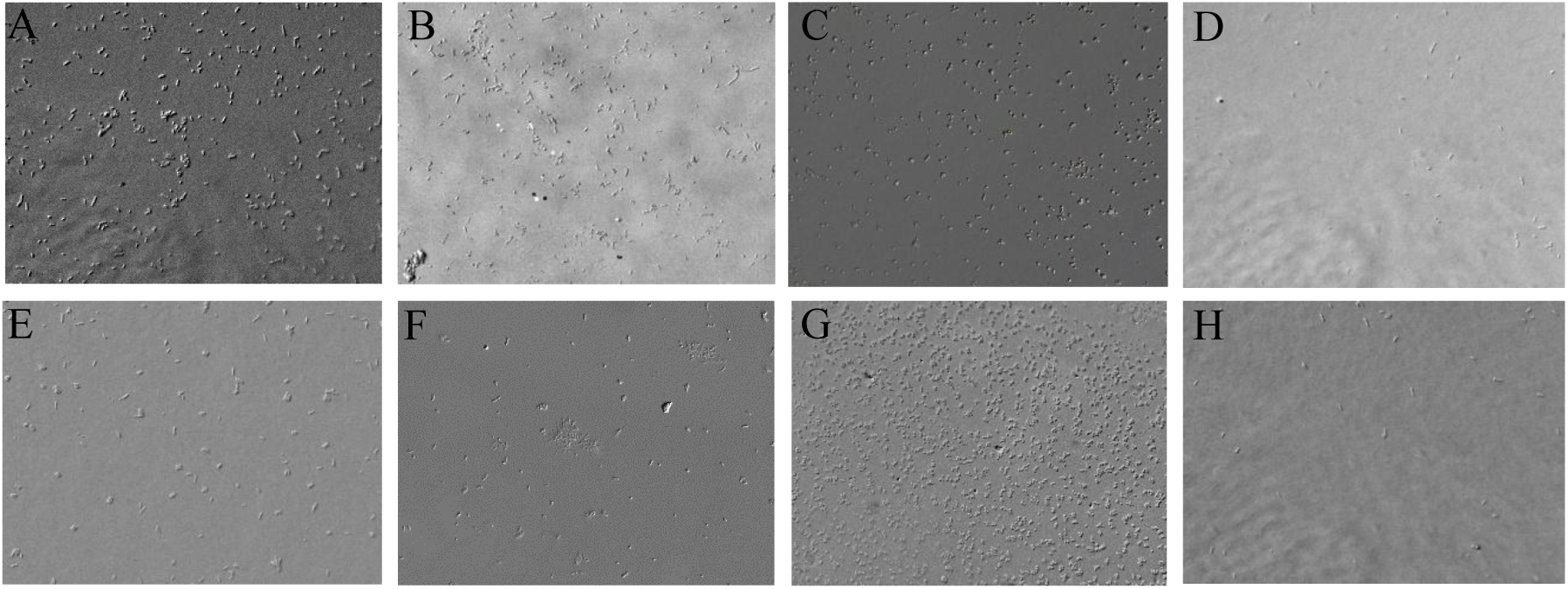
Adhesion of *P. intermedia* and *P. nigrescens*. Bright-field microscopy images acquired using a Zeiss Axioplan D2 microscope. Microscopy of biological replicates at 100× magnification and a concentration of 5×10⁷ CFU/mL. (A) adhesion of *P. intermedia* to fibronectin. (B) adhesion of *P. intermedia* to type IV collagen. (C) adhesion of *P. intermedia* to laminin. (D) negative control of *P. intermedia* (1% BSA + 1% Tween). (A) adhesion of *P. nigrescens* to fibronectin. (B) adhesion of *P. nigrescens* to type IV collagen. (C) adhesion of *P. nigrescens* to laminin. (D) negative control of *P. nigrescens* (1% BSA + 1% Tween).

### Analysis of the proteome of *P. nigrescens* adhesion to collagen type IV and laminin

As performed for the *P. intermedia* strain, after confirmation of adhesion, chromatography columns using Sepharose beads were prepared. Mass spectrometry analyses of *P. nigrescens* revealed the presence of four proteins in the type IV collagen–enriched column (Table 3), whereas five proteins were identified in the laminin type I–enriched column (Table 4). All identified proteins contained more than two unique peptides, which were exclusively associated with the identified protein group and were not found in other proteins. In addition, the protein groups did not present zero ion intensity, supporting that the identified peptides specifically belonged to the identified proteins. The proteins selected for bioinformatics analyses also showed LFQ values different from zero (Supplementary Table 1), and those with a Q-value ≤ 0.01 were considered reliable. Among the identified proteins, three were detected in both the laminin and type IV collagen column analyses, namely WP_004363718.1, EGQ15906.1, and EGQ16838.1. The presence of these three proteins in both columns highlights a likely adhesin function.

**Tabel 3:** *P. nigrescens* collagen type IV.

| First_Protein_ID | Sequence coverage [%] | Score | Q-value | Unique peptides | Coverage_class | Confidence |
| --- | --- | --- | --- | --- | --- | --- |
| WP_004363718.1 | 168 | 32331 | 0 | 4 | High | High Fidelity |
| EGQ15906.1 | 219 | 32331 | 0 | 8 | High | High Fidelity |
| EGQ16838.1 | 276 | 32331 | 0 | 5 | High | High Fidelity |
| WP_315358411.1 | 206 | 19881 | 0 | 2 | High | High Fidelity |

**Tabel 4:**
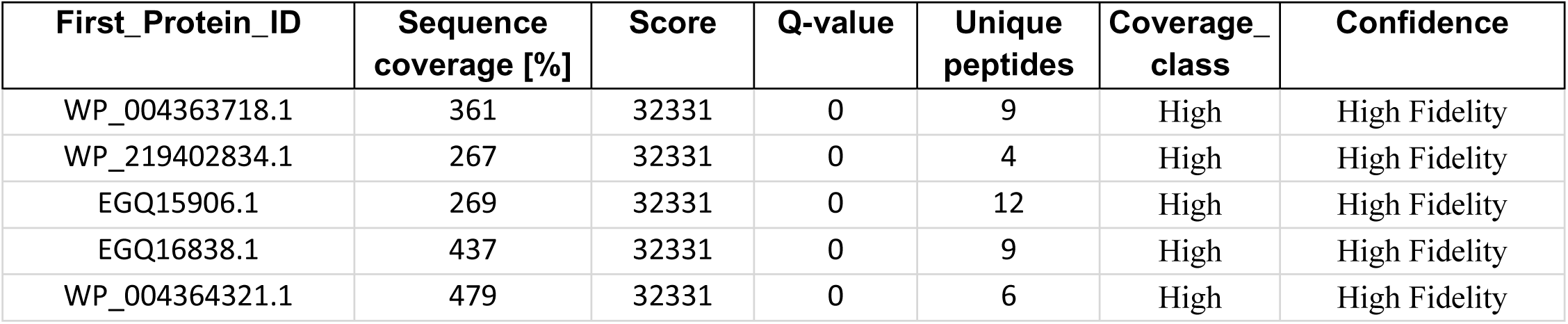
*P. nigrescens* laminin.

| First_Protein_ID | Sequence coverage [%] | Score | Q-value | Unique peptides | Coverage_class | Confidence |
| --- | --- | --- | --- | --- | --- | --- |
| WP_004363718.1 | 361 | 32331 | 0 | 9 | High | High Fidelity |
| WP_219402834.1 | 267 | 32331 | 0 | 4 | High | High Fidelity |
| EGQ15906.1 | 269 | 32331 | 0 | 12 | High | High Fidelity |
| EGQ16838.1 | 437 | 32331 | 0 | 9 | High | High Fidelity |
| WP_004364321.1 | 479 | 32331 | 0 | 6 | High | High Fidelity |

Heatmap analyses based on Z-score–normalized LFQ intensities revealed distinct patterns of protein abundance between the collagen and laminin conditions. The biological replicates showed high concordance, indicating good experimental reproducibility. Among the three recurring proteins, EGQ1506.1 exhibited above-average abundance when detected in the type IV collagen column and type I laminin column. The protein EGQ16838.1 displayed above-average abundance under both conditions, whereas WP_004363718.1 showed below-average abundance in the type IV collagen condition (Figura 2).

### Biofilm formation in the presence and absence of Matrigel

The assays performed in 96-well plates demonstrated that all strains used were capable of forming biofilm, with no substantial difference in biofilm formation among the species in the absence of matrix proteins (p > 0.1359). In the presence of these proteins, however, a significant difference in biofilm formation was observed for *P. intermedia* and *P. melaninogenica* compared with *P. nigrescens*, *F. nucleatum*, and *S. epidermidis* (p > 0.0029).

When analyzing biofilm formation for each strain individually, *P. intermedia* and *P. melaninogenica* showed reductions of 37% and 39%, respectively, in biofilm production. The mean absorbance values for *P. intermedia* with and without Matrigel were 1.51 and 2.43, respectively, while for *P. melaninogenica* they were 1.34 and 2.21, respectively. This behavior was not observed for *P. nigrescens*, for which the mean absorbance with and without Matrigel was 2.25 and 2.57, respectively. Similarly, *F. nucleatum* presented mean absorbance values of 2.27 and 2.61 with and without Matrigel, respectively. The positive control, *S. epidermidis*, did not show a difference in biofilm production in the presence or absence of ECM proteins, with mean values of 2.30 and 2.64, respectively (Figure 4).

**Fig. 3.**
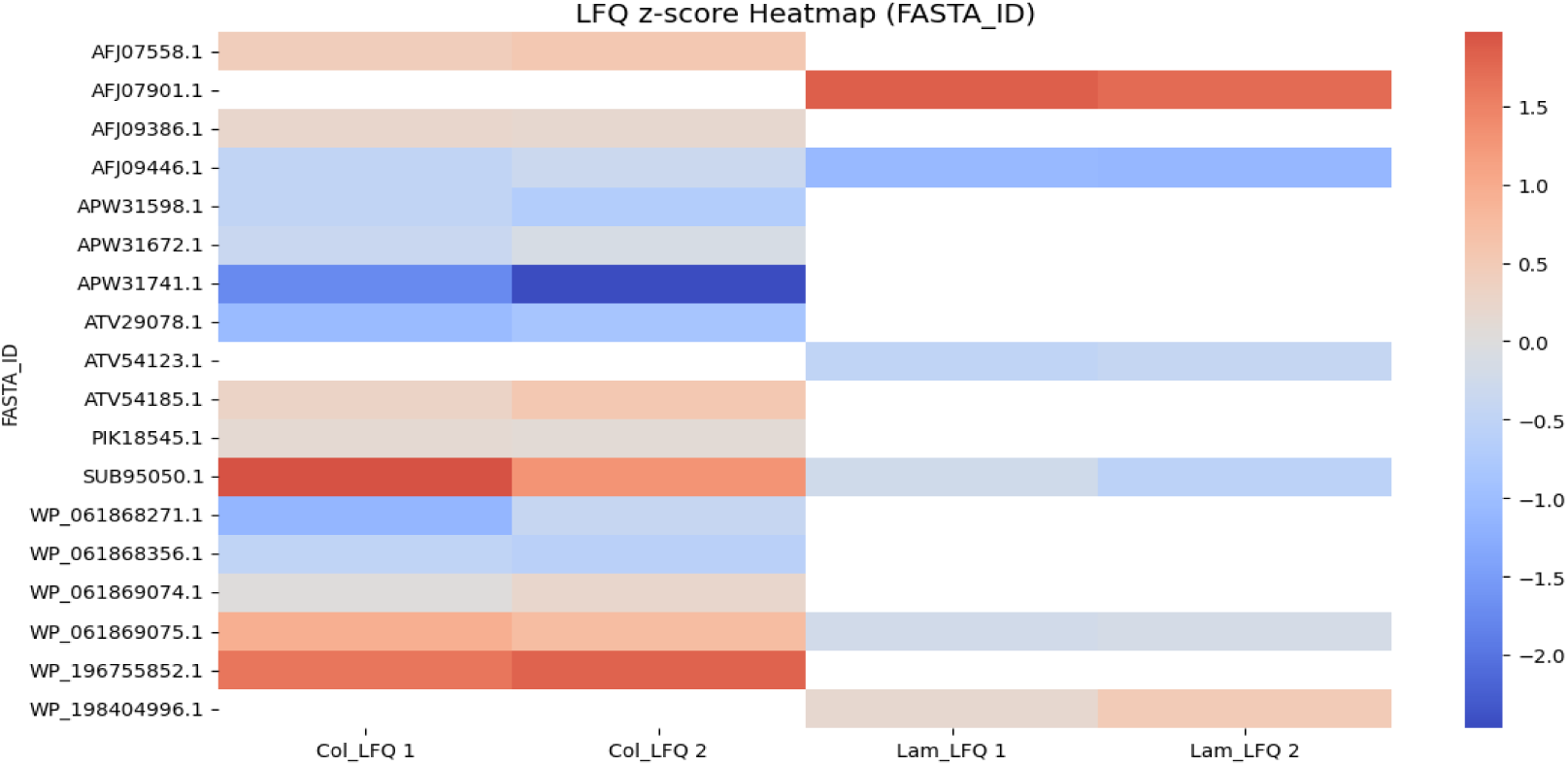
Protein abundance of *P. intermedia* identified in type IV collagen and laminin columns. LFQ Z-score heatmap of putative adhesins identified in *P. intermedia* showing the abundance of each protein across two biological replicates. Proteins with higher abundance (red) represent potential candidates for adhesin functions when compared to low-abundance proteins (blue).

**Fig. 4.**
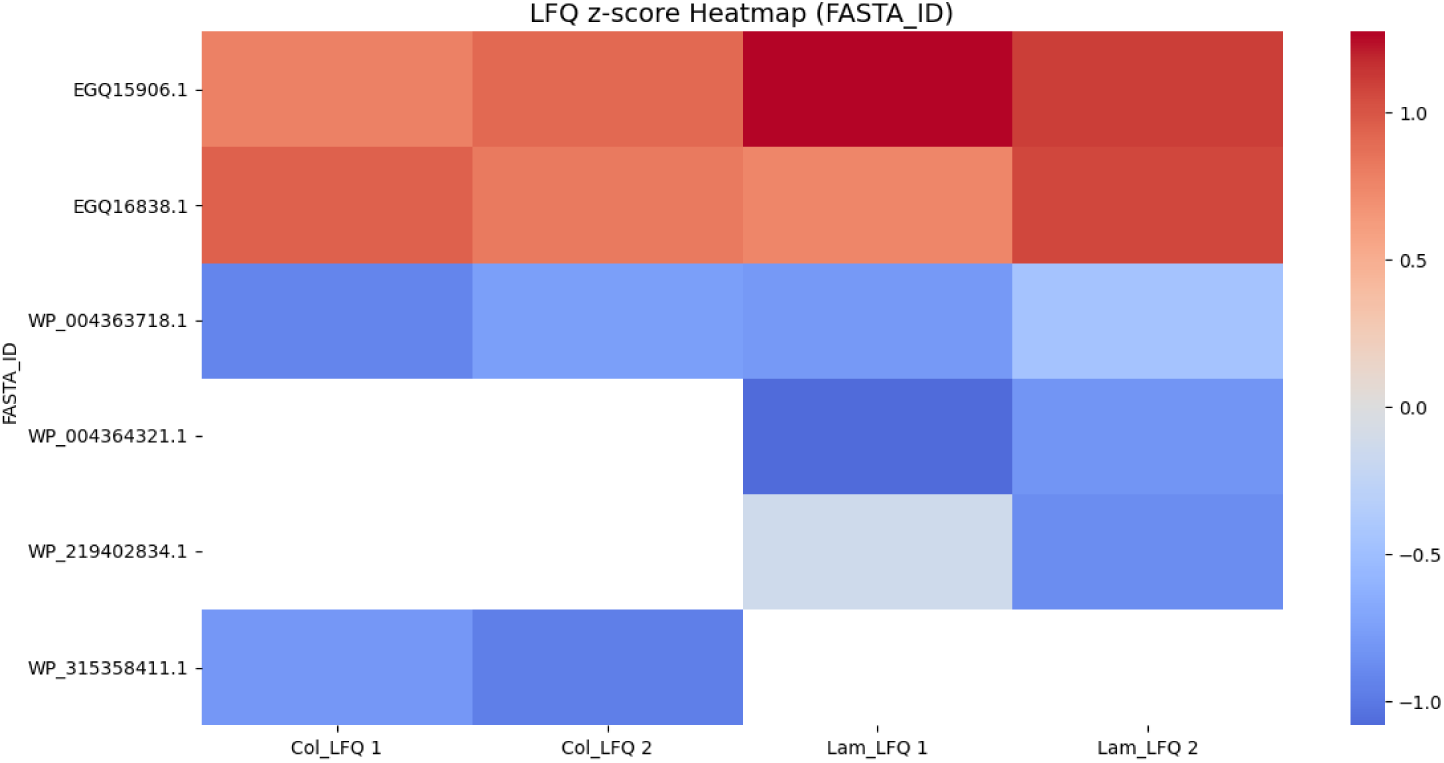
Protein abundance of *P. nigrescens* identified in type IV collagen and laminin columns. LFQ Z-score heatmap of putative adhesins identified in *P. nigrescens* showing the abundance of each protein across two biological replicates. Proteins with higher abundance (red) represent potential candidates for adhesin functions when compared to low-abundance proteins (blue).

**Figure 5:**
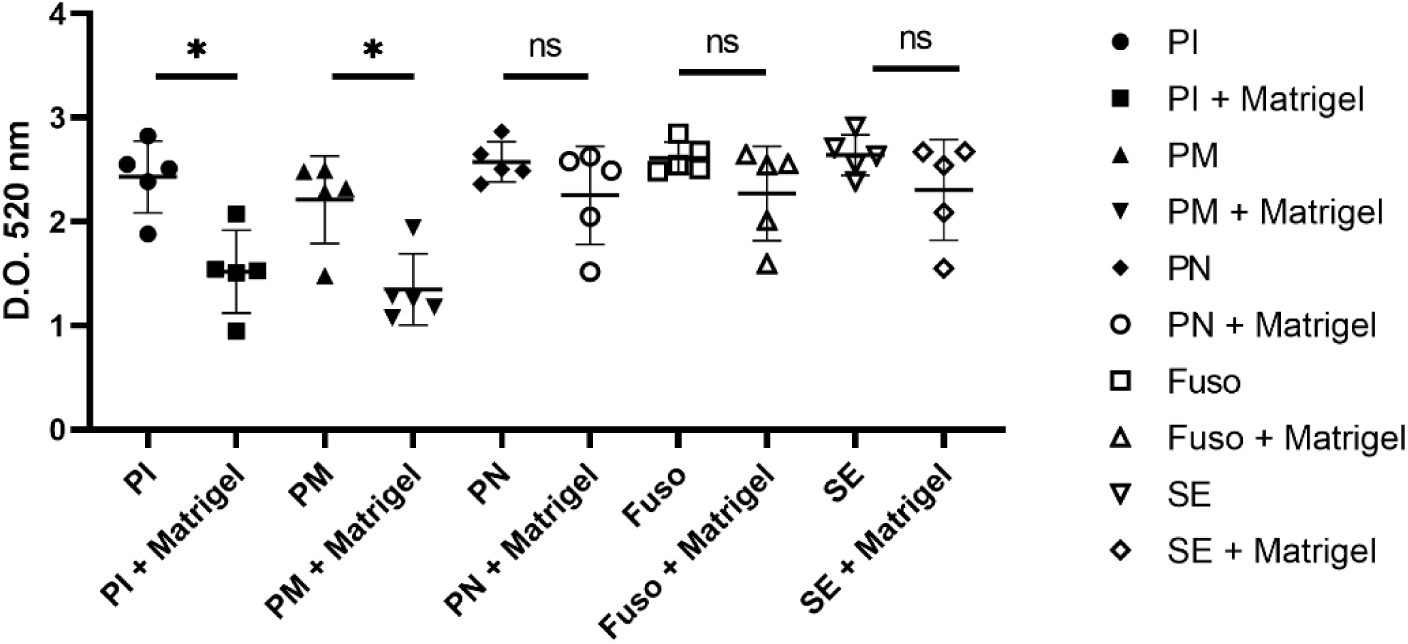
Quantification of biofilm formation in the presence and absence of Matrigel. The results represent the mean of all experiments (n = 5). Statistical analyses were performed using ANOVA followed by multiple comparisons with Tukey’s test. PI – *P. intermedia*; PM – *P. melaninogenica*; PN – *P. nigrescens*; Fuso – *F. nucleatum*; SE – *S. epidermidis*. Asterisks (*) indicate the following p values: PI vs. PI + Matrigel, p < 0.0104; PM vs. PM + Matrigel, p < 0.017.

After the initial results obtained in 96-well plates, aiming to better understand biofilm organization, analyses were performed using scanning electron microscopy. The *P. intermedia* strain, one of the species that showed differences in biofilm formation in the presence and absence of Matrigel, exhibited a distinct biofilm surface. In the absence of matrix proteins, it was possible to observe a biofilm with few bacterial aggregates (1,000× magnification) occupying the entire surface of the glass coverslip (Figure 6A). When the magnification was increased tenfold (10,000×), the presence of bacteria organizing into a biofilm and producing extracellular matrix became evident (Figure 6B).

**Figure 6:**
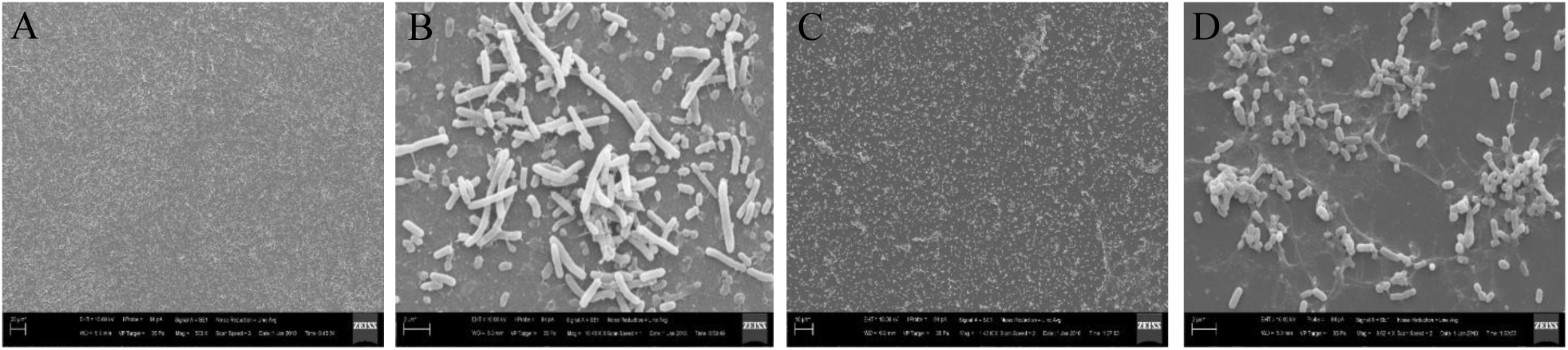
Scanning electron microscopy of *P. intermedia* biofilm in the presence and absence of Matrigel. Images were acquired using a Zeiss EVO MA10 microscope. (A) and (B) *P. intermedia* in the absence of Matrigel. (C) and (D) *P. intermedia* in the presence of Matrigel. The scale between images A, B, C, and D is 1:10. (A) 1,000× magnification, 20 µm. (B) 10,000× magnification, 2 µm.

When ECM proteins coated the bottom of the coverslip, the *P. intermedia* strain was not able to form a homogeneous biofilm, instead aggregating only in microdomains (Figures 6C and 6D). This finding corroborates the experiments performed using crystal violet staining, in which a significant reduction in biofilm production was also observed in the presence of Matrigel.

The *P. melaninogenica* biofilm exhibited a unique characteristic, since, similarly to the adhesion experiments performed on a glass surface containing ECM proteins, the bacterium was unable to adhere and form a biofilm on this type of surface (Figures 7A and 7B). A similar profile was observed in the presence of Matrigel, where few bacteria were able to adhere, although the structures of ECM proteins, such as type IV collagen, could be easily observed (Figures 7C and 7D).

**Figure 7:**
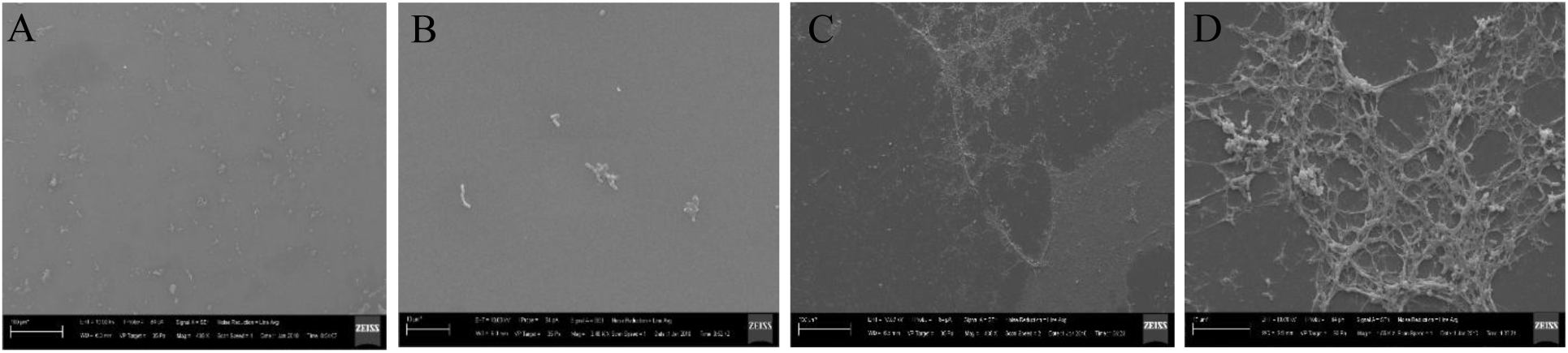
Scanning electron microscopy of *P. melaninogenica* biofilm in the presence and absence of Matrigel. Images were acquired using a Zeiss EVO MA10 microscope. (A) and (B) *P. melaninogenica* in the absence of Matrigel. (C) and (D) *P. melaninogenica* in the presence of Matrigel. The scale between images A, B, C, and D is 1:10. (A) 1,000× magnification, 20 µm. (B) 10,000× magnification, 2 µm.

As observed in the 96-well plate experiment, the *P. nigrescens* strain was able to produce biofilm both in the presence and absence of Matrigel. Visually, the biofilm appeared to be composed of dispersed cells with few aggregates, covering the entire surface of the glass coverslip. However, at higher magnification, differences in the biofilm became evident, particularly regarding cell morphology. In the absence of Matrigel, the bacterial cells were smaller (Figures 8A and 8B), whereas in the presence of ECM proteins, the cells appeared larger and more elongated (Figures 8A and 8B).

**Figure 8:**
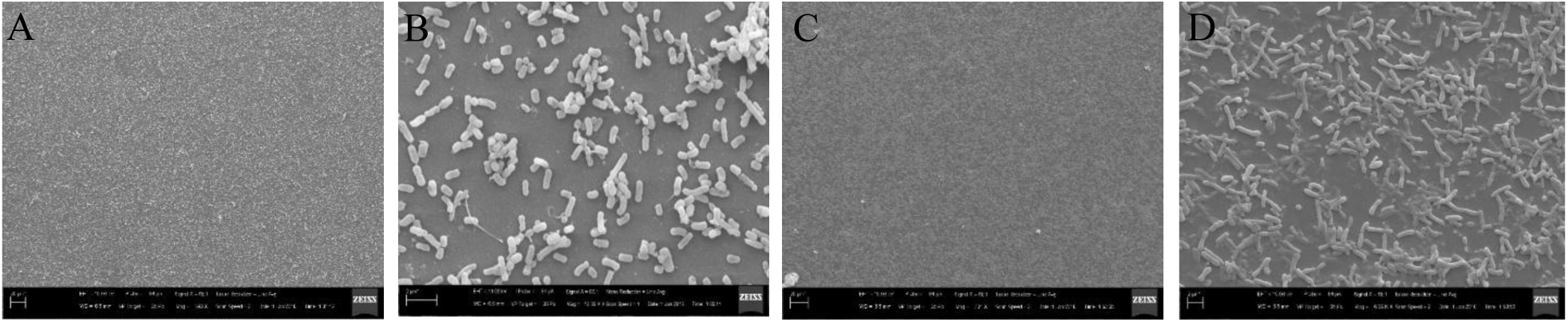
Scanning electron microscopy of *P. nigrescens* biofilm in the presence and absence of Matrigel. Images were acquired using a Zeiss EVO MA10 microscope. (A) and (B) *P. nigrescens* in the absence of Matrigel. (C) and (D) *P. nigrescens* in the presence of Matrigel. The scale between images A, B, C, and D is 1:10. (A) 1,000× magnification, 20 µm. (B) 10,000× magnification, 2 µm.

**Figure 8:**
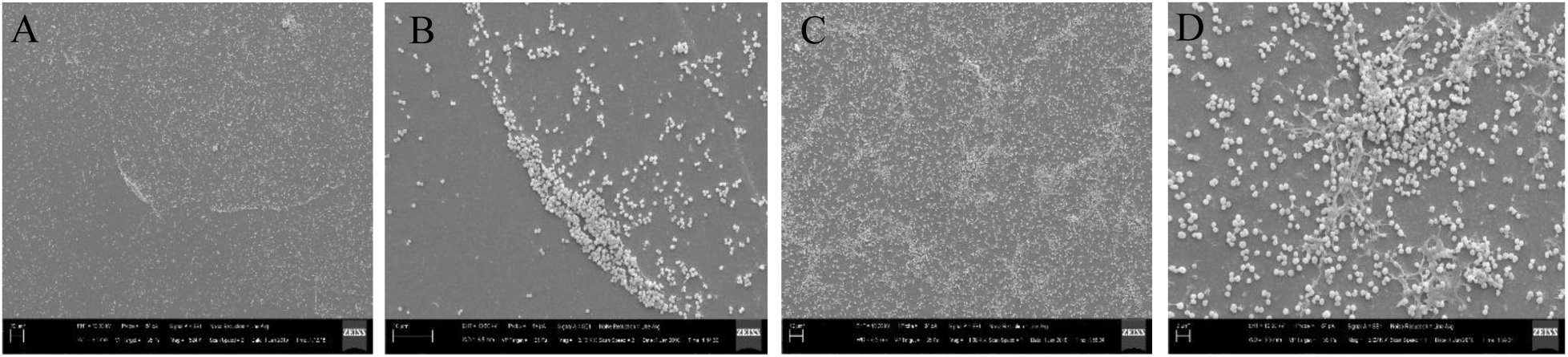
Scanning electron microscopy of *S. epidermidis* biofilm in the presence and absence of Matrigel. Images were acquired using a Zeiss EVO MA10 microscope. (A) and (B) *S. epidermidis* in the absence of Matrigel. (C) and (D) *S. epidermidis* in the presence of Matrigel. The scale between images A, B, C, and D is 1:10. (A) 1,000× magnification, 20 µm. (B) 10,000× magnification, 2 µm.

The positive control, *S. epidermidis*, demonstrated biofilm production both in the presence (Figures 24A and 24B) and absence of Matrigel, exhibiting a larger and more organized biofilm pattern in the presence of ECM molecules (Figures 25A and 25B).

### Matrigel invasion

The experiment was performed using the three *Prevotella* species. Among the three species tested, only *P. melaninogenica* was able to traverse the Matrigel layer and reach the 1× PBS buffer. *P. intermedia* and *P. nigrescens* did not demonstrate invasive capacity in any of the assays. The results were analyzed by visualizing colony growth on plates after 24, 48, and 72 hours of incubation. From the graph, it can be observed that there was no significant difference (p = 0.2427) in invasion between 48 and 72 hours (Figure 9). Analysis of the images shows an increase in *P. melaninogenica* colonies between 24 and 72 hours at the 1:100.

**Figure 9:**
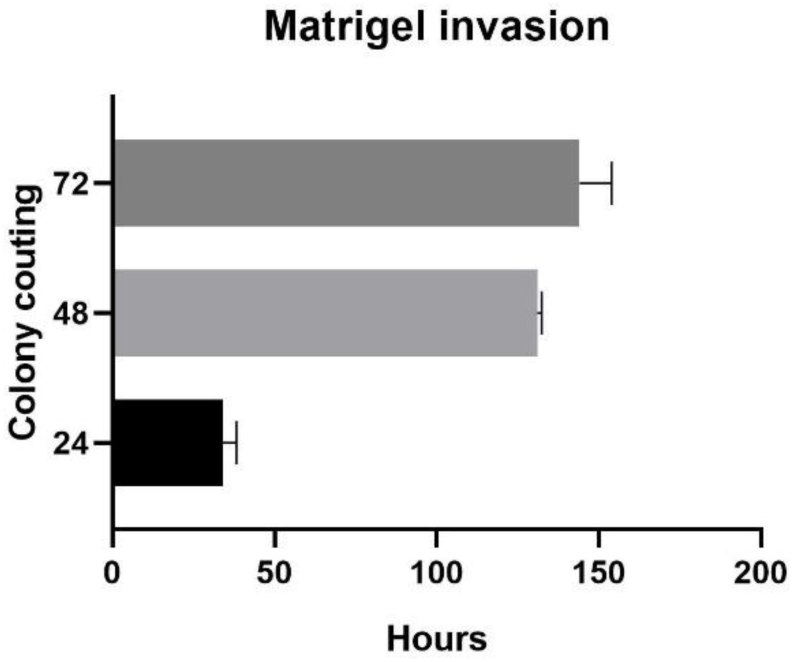
Prevotella invasion through Matrigel. Graphical representation of colony counting of *P. melaninogenica*. A volume of 10 µL was plated.

## Discussion

Bacteria that colonize the oral cavity are affected by constant environmental changes [21] and therefore require a high adhesive capacity, since feeding, mastication, tongue movements, and saliva are mechanisms that hinder bacterial colonization at these body sites. In the present study, we demonstrated that the tested species of the genera *Prevotella* and *Fusobacterium* were able to bind to ECM proteins and form biofilms in the presence of ECM. In addition, we identified that *P. melaninogenica* is capable of degrading ECM proteins and activating the plasminogen system.

The oral cavity contains ECM proteins in its composition, such as laminin, type IV collagen, fibronectin, among others. These molecules are essential for the structural support of facial tissues, including teeth, oral mucosa, palate, and gingiva [22]. In this study, the strains of *P. intermedia*, *P. nigrescens*, and *F. nucleatum* were able to adhere to laminin, type IV collagen, type I collagen, and fibronectin. The adhesive capacity of other *P. intermedia* strains has already been documented by some authors: Grenier and colleagues (1996) [23] described that the BMH strain was able to bind to type IV and type I collagen, and Marre (2020) [24] reported that an ATCC strain of *P. intermedia* is capable of binding to type I laminin; however, the adhesion mechanisms of this species have not yet been elucidated. Data obtained from the adhesion assays of *P. nigrescens* showed that this species exhibited proportionally lower adhesion than *P. intermedia*. This finding may be a result of the close association of *P. intermedia* with periodontal diseases [25], in contrast to *P. nigrescens*, which can be isolated both from infectious conditions and from individuals with oral health [15].

Adhesion to Matrigel by *P. intermedia* and *P. nigrescens* was previously documented by Alugupalli and Kalfas in 1997 [26]; however, as in the present study, adhesion was not quantitatively assessed. Our work suggests that the difficulty in quantifying adhesion is due to the presence of bacterial aggregates. Co-aggregation of bacterial species is very common in the oral cavity, where several microorganisms are able to bind to each other, forming a symbiotic biofilm [27], which protects them from external and internal agents such as tooth brushing and swallowing. To elucidate the adhesion mechanisms of the three species, laminin [24] and type IV collagen were selected to identify possible adhesins of *P. intermedia* and *P. nigrescens*, due to their high adhesion rates. For *F. nucleatum*, fibronectin and type I collagen were tested. It is important to note that pilot tests were performed with type IV collagen and *F. nucleatum*; however, adhesion quantification was complex, since the bacterium formed aggregates around the collagen fibers. With this in mind, the study proceeded by investigating adhesion only to type I collagen.

In recent years, only one study (Yu et al., 2006) was able to identify an adhesin, AdpB, from *P. intermedia* capable of binding to fibronectin. Based on this, we used a proteomics approach through mass spectrometry to identify potential adhesive molecules. We identified a total of six possible laminin adhesins, including two OmpA proteins, PG33 (ATV54123.1) and PG32 (WP_198404996.1), which contain adhesion domains (IPR006665) and a type VI secretion system domain (PF00691). The type VI secretion system is a virulence factor of Gram-negative bacteria (Coulthurst, 2019) and is important for the translocation of effector proteins to the surface of host cells. Notably, this pathogenicity mechanism has already been associated with adhesion of *Klebsiella pneumoniae* strains during gastrointestinal tract colonization [28] and with adhesion of avian *E. coli* [29].

The two TonB proteins identified (AFJ09446.1; WP_061869075.1) do not contain specific adhesion domains. TonB proteins are essential for iron transport in Gram-negative bacteria, binding to siderophores, nickel complexes, vitamin B12, and carbohydrates [30]. Their ability to bind carbohydrates is particularly interesting, since proteins such as collagen, laminin, and fibronectin are glycosylated and contain multiple carbohydrate-binding domains. Pauer et al. (2009) [31] demonstrated that a TonB protein from *B. fragilis* was responsible for adhesion of this microorganism to fibronectin, supporting the identification of TonB as a potential adhesin. The SurA protein (AFJ07901.1) also plays an essential role in the adhesion of uropathogenic *E. coli* (UPEC) to bladder cells [32] and may therefore be another possible adhesin candidate. Together, these findings and our results indicate that some proteins with adhesin function are moonlighting proteins, that is, proteins with multiple functions. The uncharacterized protein (SUB95050.1) did not show a function directly associated with adhesion but is located in the outer membrane and may also act as an adhesin.

Proteomic analysis of adhesins binding to type IV collagen identified 15 possible adhesins, among which three—OmpA (WP_198404996.1), TonB (AFJ09446.1), and the uncharacterized protein (SUB95050.1)—had already been identified in the laminin affinity column. The ability of a single bacterial molecule to bind to multiple ECM components has been previously described. In 1989, Emtidy et al. showed that the YadA protein of *Yersinia* was capable of binding to laminin, fibronectin, and collagen. In 2007, the FnBPA protein of *Staphylococcus aureus* was described as an adhesin capable of binding to fibronectin and elastin [33]. Adhesins with the ability to bind more than one ECM protein are known as MMSCRAMs. To date, no MMSCRAMs had been described in species of the genus *Prevotella*.

Other potential adhesins were also identified, including two TonB proteins (AFJ07558.1; WP_061869075.1), an OmpA protein (PIK18545.1), and the RagB/SusD protein (WP_061869074.1), which contains a binding domain associated with sugar metabolism–related proteins, especially in oral pathogens such as *P. gingivalis* and *T. forsythia* [37]. RagB/SusD is an outer membrane protein strongly associated with virulence in these bacterial species. Considering that ECM proteins contain sugars in their composition, it is reasonable to suggest that the identified RagB/SusD protein may function as an adhesin in *P. intermedia*. Some hypothetical proteins (APW31598.1; APW31672.1; ATV29078.1) were also identified. Although none possess domains directly related to adhesion, their porin domains may be involved in ECM adhesion. Paulson et al. (2015) reported that a porin from *P. aeruginosa* is essential for bacterial virulence in cystic fibrosis, as this molecule is able to bind to vitronectin. The same may be inferred for the outer membrane beta-barrel protein (WP_061868271.1), which also contains a porin domain.

In addition to the proteomic results obtained for *P. intermedia*, we were also able to identify potential adhesins of *P. nigrescens*. Analysis of laminin-binding molecules revealed the presence of five proteins, including two hypothetical proteins (EGQ15906.1; EGQ16838.1), one OmpA protein (WP_004363718.1), and one copper resistance protein (WP_219402834.1), all of which contain domains involved in protein binding and porin function and are localized in the outer membrane. Finally, a protein associated with cell division (WP_004364321.1) was identified, although it has not been described as an adhesin. As observed for *P. intermedia*, the proteins identified as collagen adhesins in *P. nigrescens* had already been identified as laminin adhesins, further supporting the idea that both species may possess MMSCRAMs. However, unlike *P. intermedia*, no adhesins of *P. nigrescens* had been previously identified, highlighting the importance of our work in elucidating the initial colonization process of these species.

As previously stated, *P. melaninogenica* was the only species that did not exhibit adhesion potential in the assays with type IV collagen, fibronectin, and Matrigel. To date, there are few studies relating this species to adhesion to ECM proteins or demonstrating its association with other species. This bacterium has been isolated from sputum samples of children with cystic fibrosis [34] and from patients with advanced periodontitis [35]. Despite these reports, *P. melaninogenica* is considered part of a healthy microbiota and has been less frequently associated with disease when compared with the other species selected in this study [15; 36; 38]. Based on our results, we believe that *P. melaninogenica* may harbor a higher number of virulence factors with aggressive potential, such as proteolytic enzymes and plasminogen activation, among others, and fewer adhesion factors to ECM proteins, which could explain the lack of adhesion to these molecules under the tested conditions.

Adhesion is a crucial step for biofilm formation [39], and Blanco-Romero et al. (2024) [40] described that the presence of ECM proteins is essential for biofilm formation by *Pseudomonas ogarae*. With this in mind, we investigated biofilm development in the presence and absence of Matrigel. Our results from the 96-well plate assays demonstrated that all species were able to produce biofilm both in the presence and absence of Matrigel. However, *P. intermedia* and *P. melaninogenica* showed a significant reduction in biofilm production in the presence of Matrigel. This result was surprising, since studies evaluating biofilm formation in the presence of matrix proteins [40;41] have shown that these molecules promote biofilm structuring. When living in biofilms, bacteria produce a polymeric matrix that functions in protection, nutrient acquisition, and even communication. Our hypothesis is that the presence of Matrigel may act as a substitute for the polymeric matrix produced by bacteria, as its structure is rich in carbohydrates and structural proteins that provide protection and nutrients. Thus, *P. intermedia* and *P. melaninogenica* exhibited a reduction in biomass attributed to their own biofilm matrix.

Based on these results, we sought to further understand why *P. intermedia* and *P. melaninogenica* showed reduced biofilm production in the presence of Matrigel. To this end, we performed a biofilm formation assay in 24-well plates with glass coverslips, followed by scanning electron microscopy analysis. This approach allowed us to observe the biofilm surface and assess how these species reorganize in the presence of Matrigel.

Our results demonstrated that the *P. intermedia* strain indeed exhibits a different distribution on the biofilm surface in the presence of Matrigel, with cells adhering only where ECM molecules are present and failing to form a classical biofilm. In contrast, *P. melaninogenica* did not show biofilm formation either in the presence or absence of Matrigel, reinforcing the adhesion assay results, in which this microorganism was unable to initiate adhesion on glass surfaces. The species *P. nigrescens* exhibited biofilm formation with greater aggregation on Matrigel-coated slides, corroborating studies showing that this species is capable of forming strong and dense biofilms [42].

With adhesion and biofilm formation confirmed, the next step was to deepen our understanding of the interactions between these oral pathogens and the ECM. To this end, we performed a tissue invasion model using a Matrigel layer in Transwell plates. This type of assay is widely used to study the translocation of cancer cells [43]; however, Korhonen and colleagues (1992) [44] tested Matrigel invasion by *E. coli*, a bacterium with invasive capacity. Following this work, several studies have continued to investigate whether bacterial pathogens are able to invade the basement membrane [45]. Surprisingly, in our study, the only species capable of crossing the Matrigel layer was *P. melaninogenica*, precisely the strain that was unable to adhere but was nevertheless able to translocate in the Transwell system. In 2022, a study by Zheng et al. [46] demonstrated that this species is capable of invading epithelial cells and the lamina propria under conditions of prior mucosal inflammation. Our work shows that this *P. melaninogenica* strain is able to translocate across the ECM layer independently of a pre-existing disease condition.

The Matrigel used is composed of several matrix molecules, such as laminin, type IV collagen, and even plasminogen [47]. Based on the invasion model results, we then decided to test the bacterial capacity to degrade Matrigel and plasminogen. Our results indicated that *P. melaninogenica* was the only species capable of degrading the proteins present in Matrigel and the purified plasminogen molecule. Zheng et al. (2022) [46] demonstrated that *P. melaninogenica* was able to translocate through Matrigel but did not describe the mechanism by which the microorganism crossed the ECM protein layer. Our results suggest that the translocation mechanism may be associated with the ability of this *Prevotella* species to activate the plasminogen system, promoting cleavage of plasminogen present in Matrigel and, consequently, disruption of this matrix.

Only the experiments with *P. melaninogenica* resulted in significant plasminogen activation rates when compared with those obtained for the positive control, plasmin. This type of virulence mechanism is well described in several aerobic species, such as streptokinase from *Streptococcus*, staphylokinase from *Staphylococcus*, and the flagellum of *E. coli*, among others [45; 48; 49]. In anaerobic bacteria, a clinical strain of *P. intermedia* has already been described as a bacterium capable of altering the plasminogen regulatory system [50]; however, there are no descriptions of this virulence mechanism in *P. melaninogenica*.

